# Reward priming differentially modulates enhancement and inhibition in auditory decision-making

**DOI:** 10.1101/2021.12.23.473984

**Authors:** Hidekazu Nagamura, Hiroshi Onishi, Momoko Hishitani, Shota Murai, Yuma Osako, Kohta I. Kobayasi

## Abstract

In cognitive sciences, rewards, such as money and food, play a fundamental role in individuals’ daily lives and well-being. Moreover, rewards that are irrelevant to the task alter individuals’ behavior. However, it is unclear whether explicit knowledge of reward irrelevancy has an impact on reward priming enhancements and inhibition. In this study, an auditory change-detection task with task-irrelevant rewards was introduced. The participants were informed explicitly in advance that the rewards would be given randomly. The results revealed that while inhibition related to reward priming only occurred when the participants were explicitly informed about rewards, implicit instruction thereof resulted in enhancement and inhibition associated with reward priming. These findings highlight the contribution of explicit information about rewards associated with auditory decisions.

## 1. Introduction

Rewards have a considerable impact on individuals’ daily lives. For instance, performances typically improve when individuals are rewarded substantially for good work. Therefore, it is imperative to address the underlying reward mechanisms that help individuals in performing efficiently.

Individuals’ behavior is influenced when rewards are not associated with the current task. For instance, when stimuli that were previously associated with rewards but are currently independent are presented, participants’ responses have been noted to be slower [1–5]. Moreover, Hickey et al. [6] revealed that a stimulus presented with a reward in the previous trial distorted the attentional process while rewards were independent of stimulus and actual performance. This previous reward bias, that is, reward priming, is believed to comprise a probabilistic association between the stimulus and reward [7].

However, the causes of the reward priming remain very ambiguous. In previous studies, participants have not always been informed that the reward was not associated with the current task goal. Therefore, it has been unclear whether reward priming is driven by explicit reward information. Numerous studies revealed that rewards given randomly that the participants knew about had different effects in comparison to contingent or performance-independent rewards in cognitive control and learning [8–11]. Information related to reward randomness may be an important aspect of reward priming.

Accordingly, the purpose of this study was to investigate the effects of reward priming on auditory decisions. To elucidate explicit information effects on reward priming, participants were informed that they would be rewarded randomly for correct answers. In addition, whether reward priming applied to auditory tasks and perceptual decision-making was investigated.

## 2. METHODS

### 2.1 Participants

The participants comprised 16 healthy students (8 females, 22–26 years old) who all provided informed consent. They all had normal or corrected-to-normal vision and normal hearing. The study was approved by the ethics board of Doshisha University.

### 2.2 Stimuli

The stimulus was a 2300–3000 ms auditory stimulus that comprised two tone burst sequences and white noise (Fig. 1A). The frequencies of each tone burst sequence (Tone A, B) were 1000 and 500 Hz, with stimulus lengths of 155 and 99 ms and inter-stimulus time intervals of 93 and 74 ms. These sequences were employed to simulate the time structure of natural sounds [12]. While the maximum sound pressure level of the stimulus was 74 dB SPL, the rise and fall of the tone bursts were both set to 5 ms. Matlab (MathWorks, Inc., Natick, MA, USA) was employed to process the auditory stimuli. The timing of the *disappearance* of the experimental stimulus was defined as the offset of the inter-stimulus interval immediately following the disappearance of the tone burst sequence. The onset of the next tone burst was expected to be the earliest to detect the disappearance of the tone burst sequence. The white noise disappeared 1500 ms after the tone burst sequence disappeared.

**Fig. 1.**
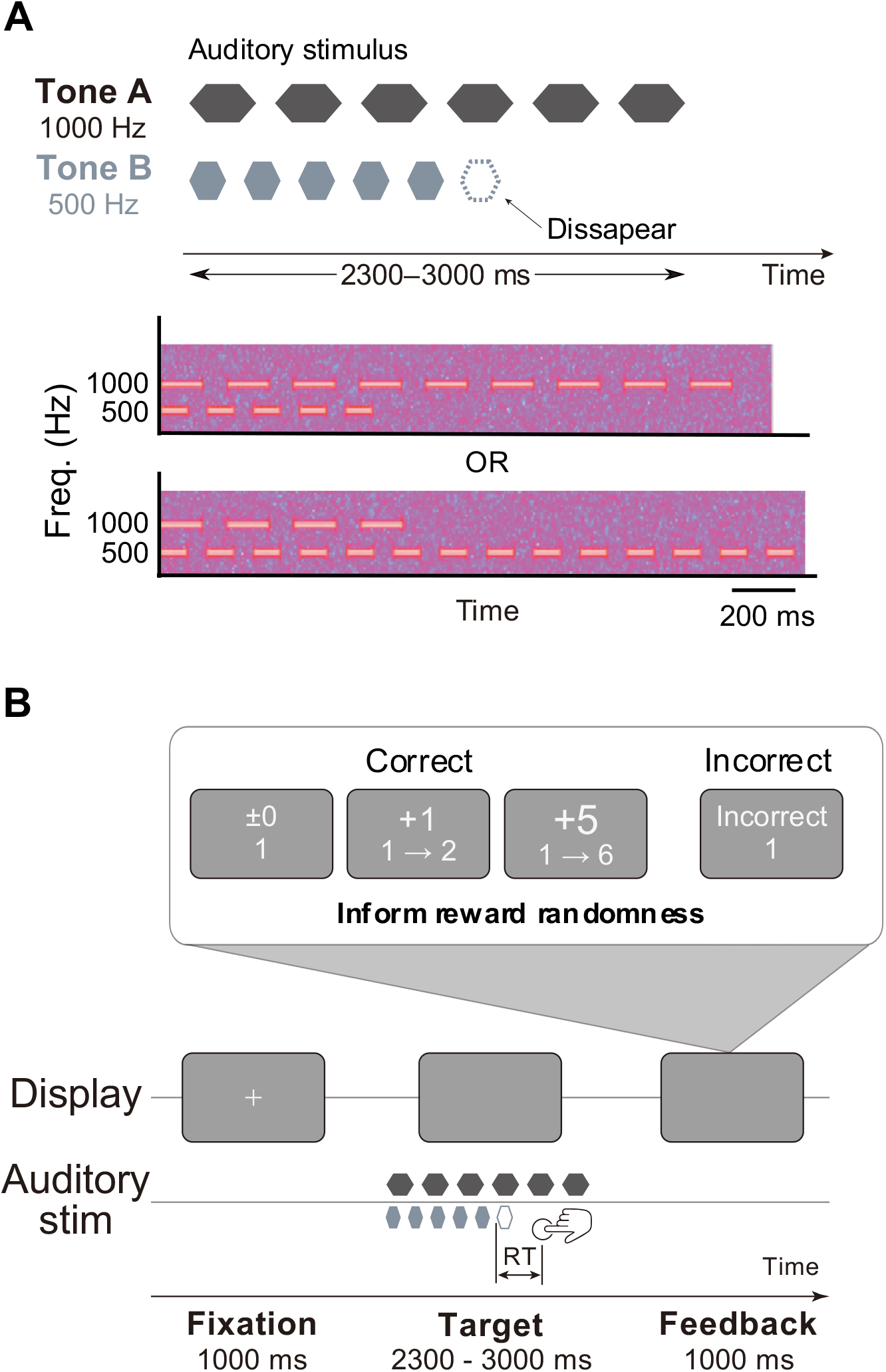
(A) The structure of sound stimuli and spectrogram. (B) The structure of task paradigm.

### 2.3 Procedure

The auditory change detection task was employed to create a unique two-alternative forced-choice (2AFC) task in order to examine the effect of reward priming (Fig. 1B). The participants were first required to look at a gazing point that was displayed at the center of the screen. After presenting the gazing point for 1000 ms, the experimental stimulus was presented. Subsequently, between 800 and 1500 ms after the presentation, either Tone A or B disappeared (Fig. 1A). The participants were then asked to as soon as possible answer which tone had disappeared. The reaction time was defined as 1500 ms after the disappearance of the presented stimulus. Accordingly, any response before the disappearance of the stimulus or outside the reaction time was deemed an incorrect answer. If the participants answered correctly, they earned 0, 1, or 5 points randomly. One of these numbers and the change of total score were displayed at the center of the screen. If the participants answered incorrectly, they did not earn any points. The word *Incorrect* and the total score were displayed at the center of the screen. Finally, the participants were rewarded in accordance with their total score. Practice consisted of 60 trials without scoring and 12 trials with scoring, followed by three sessions of 120 trials in the test experiment. The Presentation software package (Neurobehavioral Systems, Inc., Albany, CA, USA) was employed to program the experiment.

### 2.4 Analysis

To exclude inter-participant variability, data for individual trials were converted to z-scores for each participant. To examine the association between stimulus and reward, reaction time was analyzed by determining whether the stimulus was the same as the previous one (Fig. 2). The details thereof are discussed in the Results section. The statistical analyses of behavioral data for Kendall correlation test and paired t-test were conducted in Python with the package Pingouin [13].

**Fig. 2.**
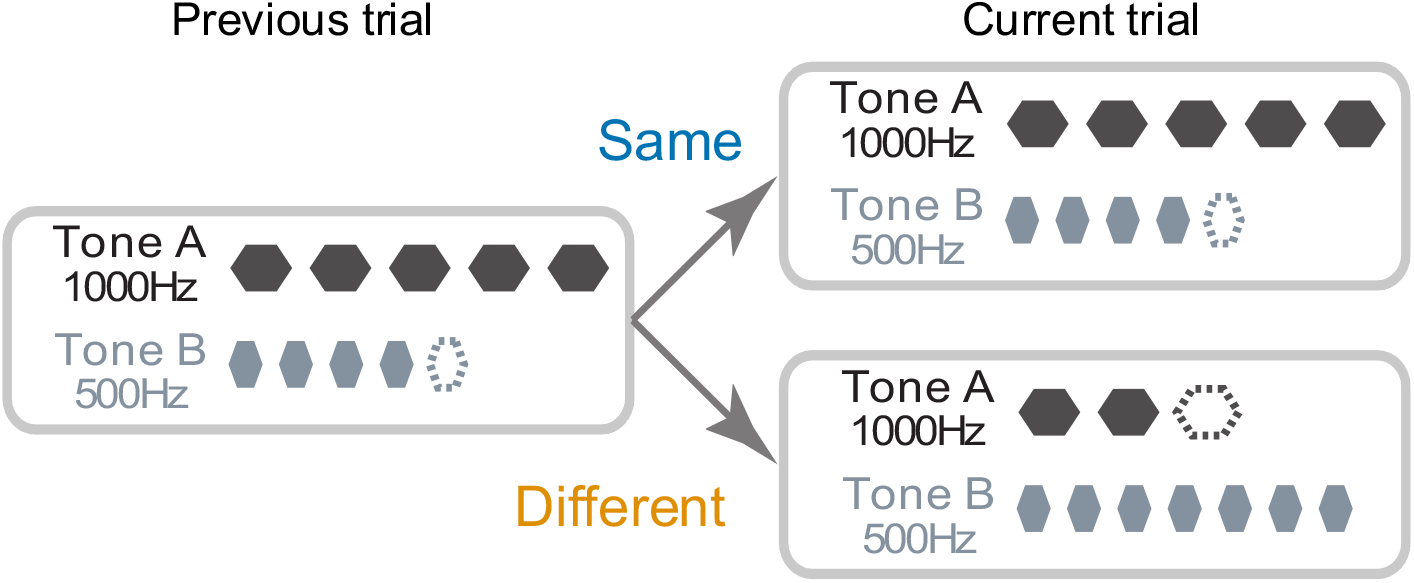
The conceptual schematics of previous-current stimulus condition.

## 3 RESULTS

All the participants (n = 16) performed the task well (Fig. 3) and their reaction times were not influenced by the current reward (*r* = 0.011, *p* > 0.1, Kendall correlation test; Fig. 4) because the rewards were not associated with stimuli. On the contrary, when the previous trial reward had a high value, the participants were slower (*r* = 0.38, *p* < 0.001, Kendall correlation test; Fig. 5), thereby revealing that reward priming influences decisions even when the participants were informed explicitly that the rewards were unrelated to the task. The effects of both previous and the current trial stimuli were tested (Fig. 2). The results revealed that their reaction times were slower when the stimulus was the same as the previous trial than when it was different (*t*(15) = 4.17, *p* < 0.001, paired t-test; Fig. 6). This is contradictory to previous results like sequential effect [14]. This strange phenomenon may be caused by the experiment design, such as detecting the disappearance of the stimulus and tone burst sequences. Moreover, when the current stimulus differed from the previous one, based on the extent of the previous reward, the response speeds were slower. However, when the current stimulus was the same as the previous one, there was no difference in reaction times in comparison to the previous reward size (*r* = 0.18, *p* > 0.1, and *r* = 0.38, *p* < 0.01, respectively, Kendall correlation test with Bonferroni correction; Fig. 7 blue and orange line).

**Fig. 3.**
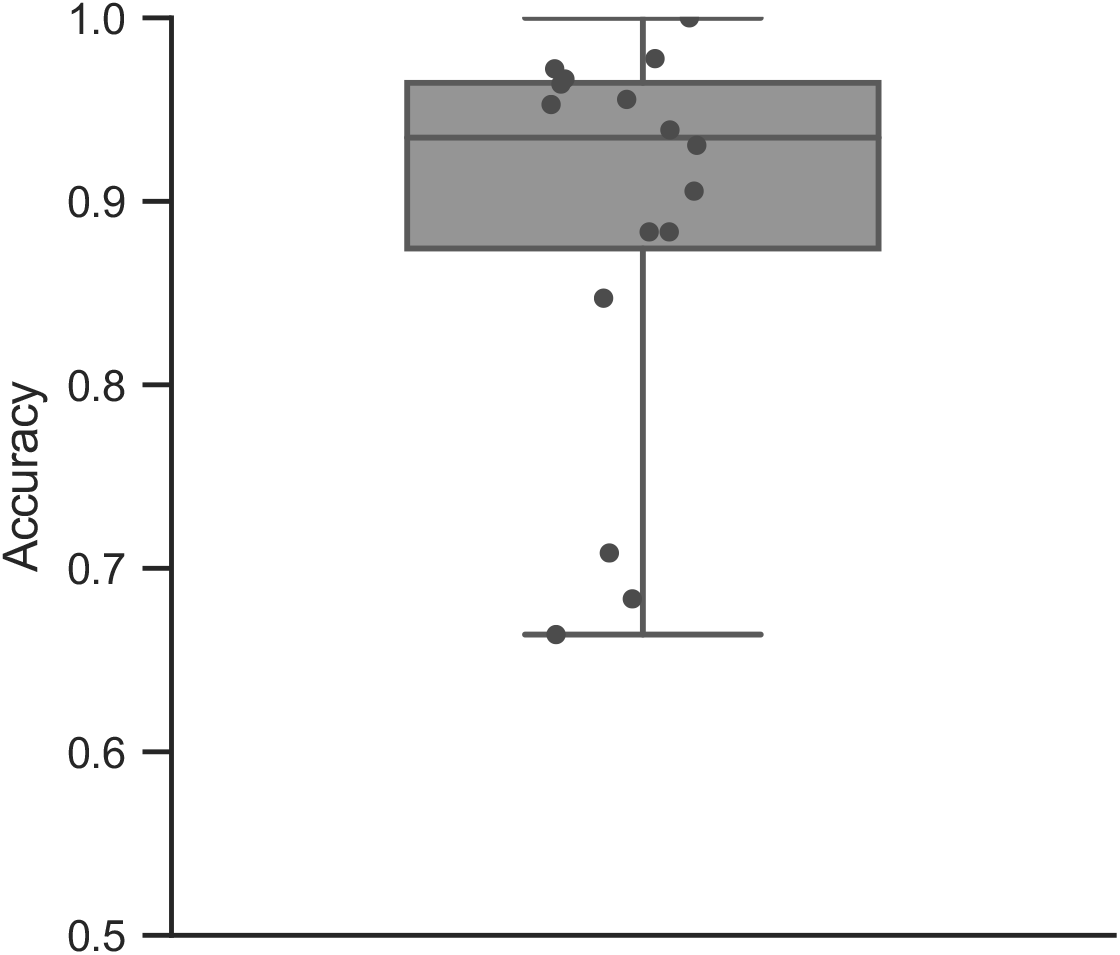
The boxplot of participant accuracy. Each point represents the accuracy for individual participants.

**Fig. 4.**
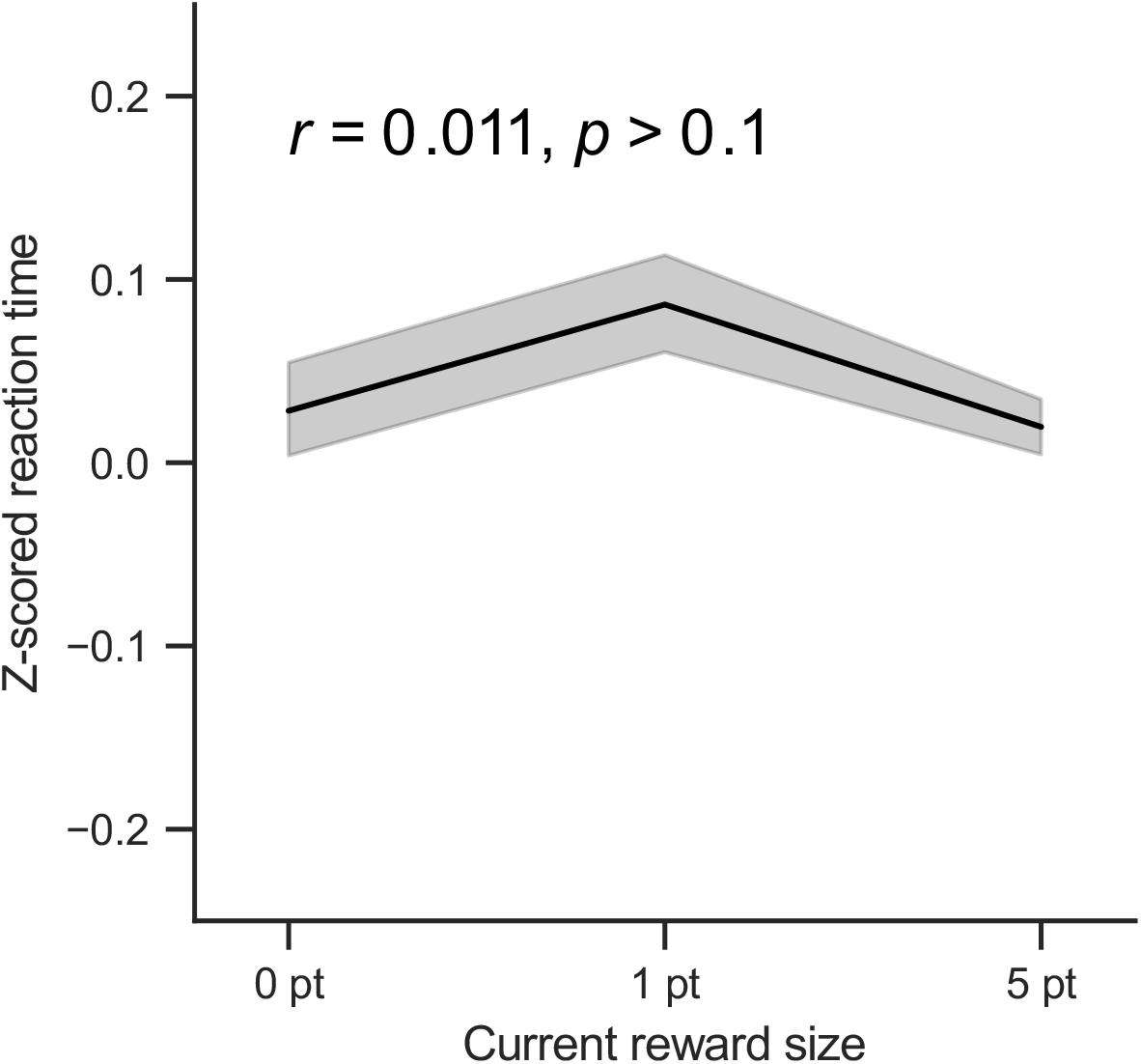
Mean z-scored reaction time (RT) in current reward size. Shaded areas denote standard error of mean (SEM).

**Fig. 5.**
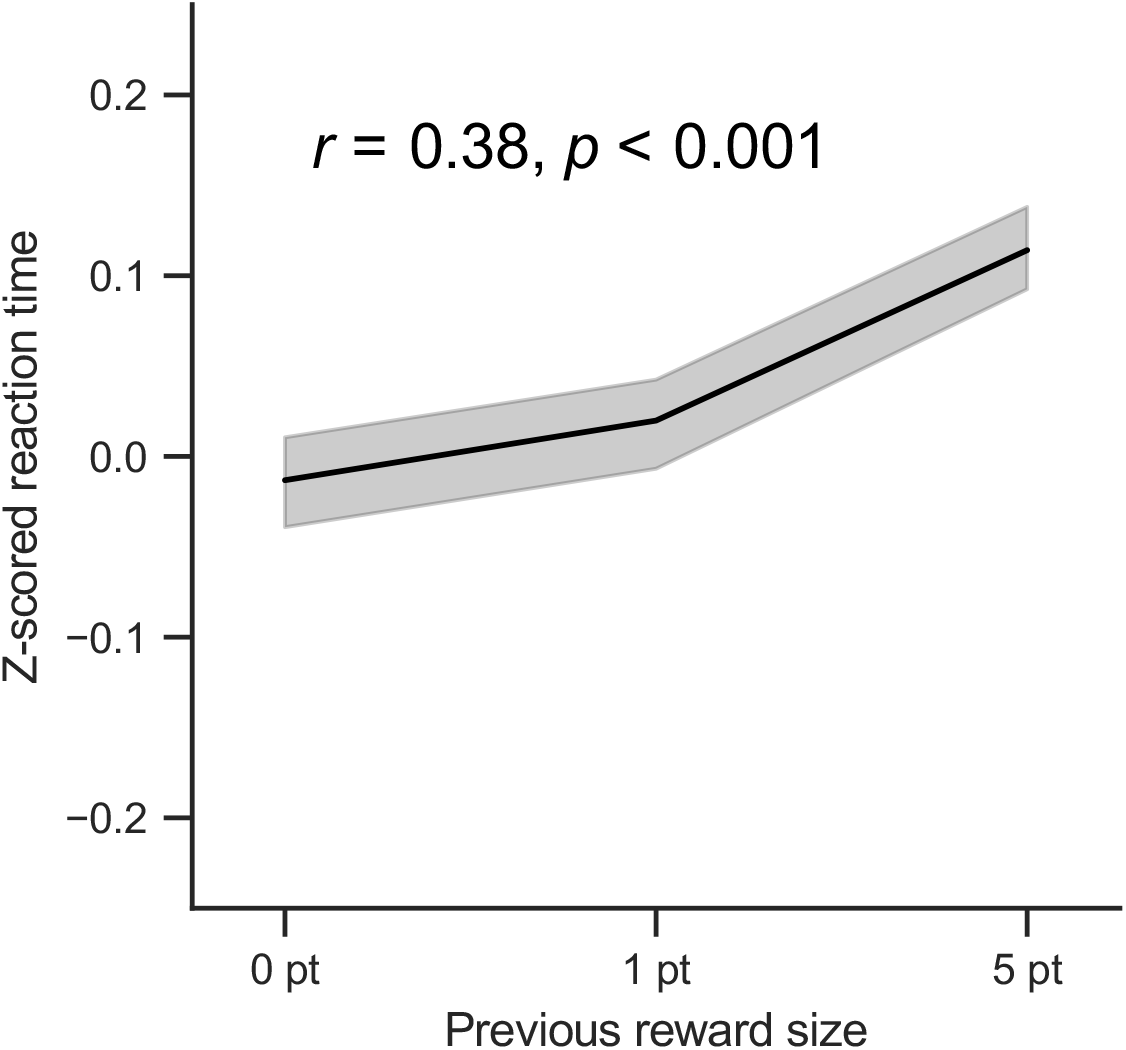
Mean z-scored RT in previous reward size.

**Fig. 6.**
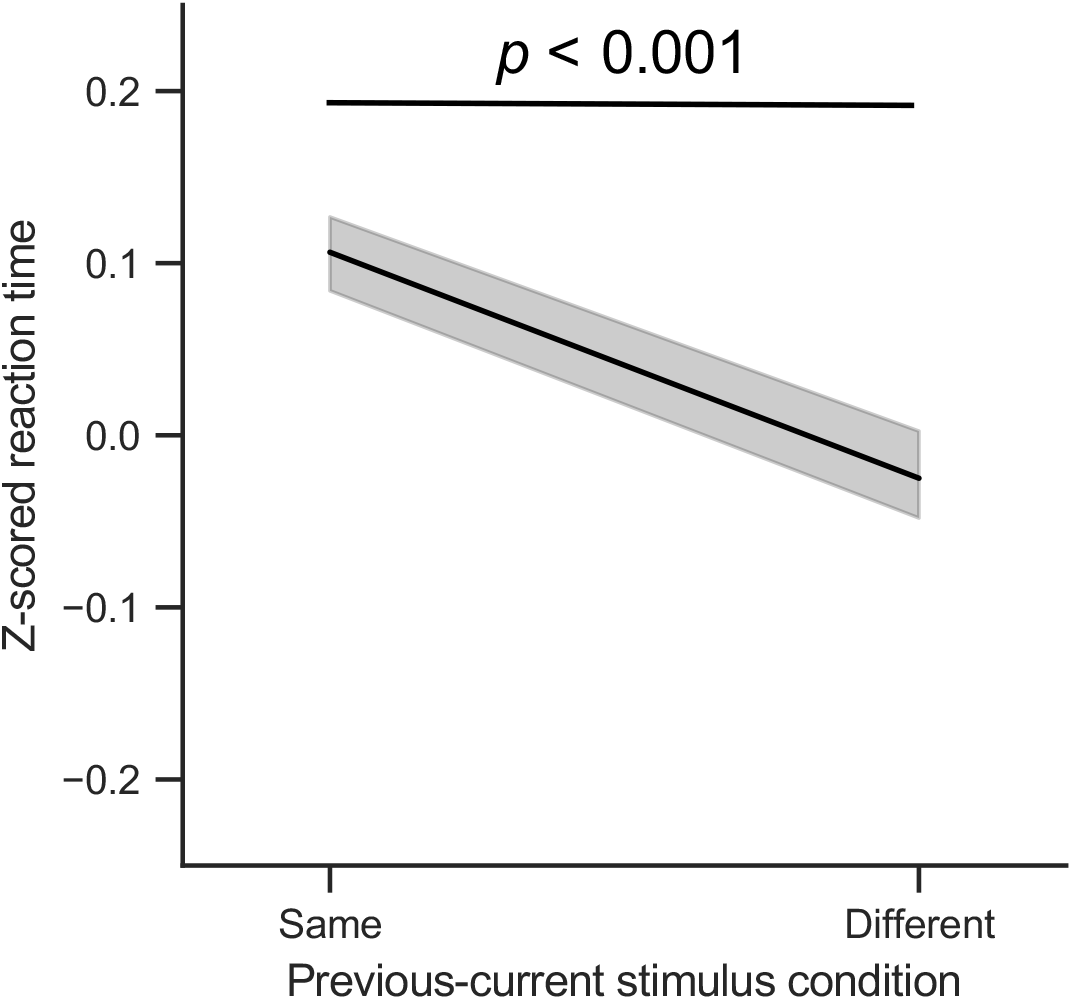
Mean z-scored RT in previous-current stimulus condition.

**Fig. 7.**
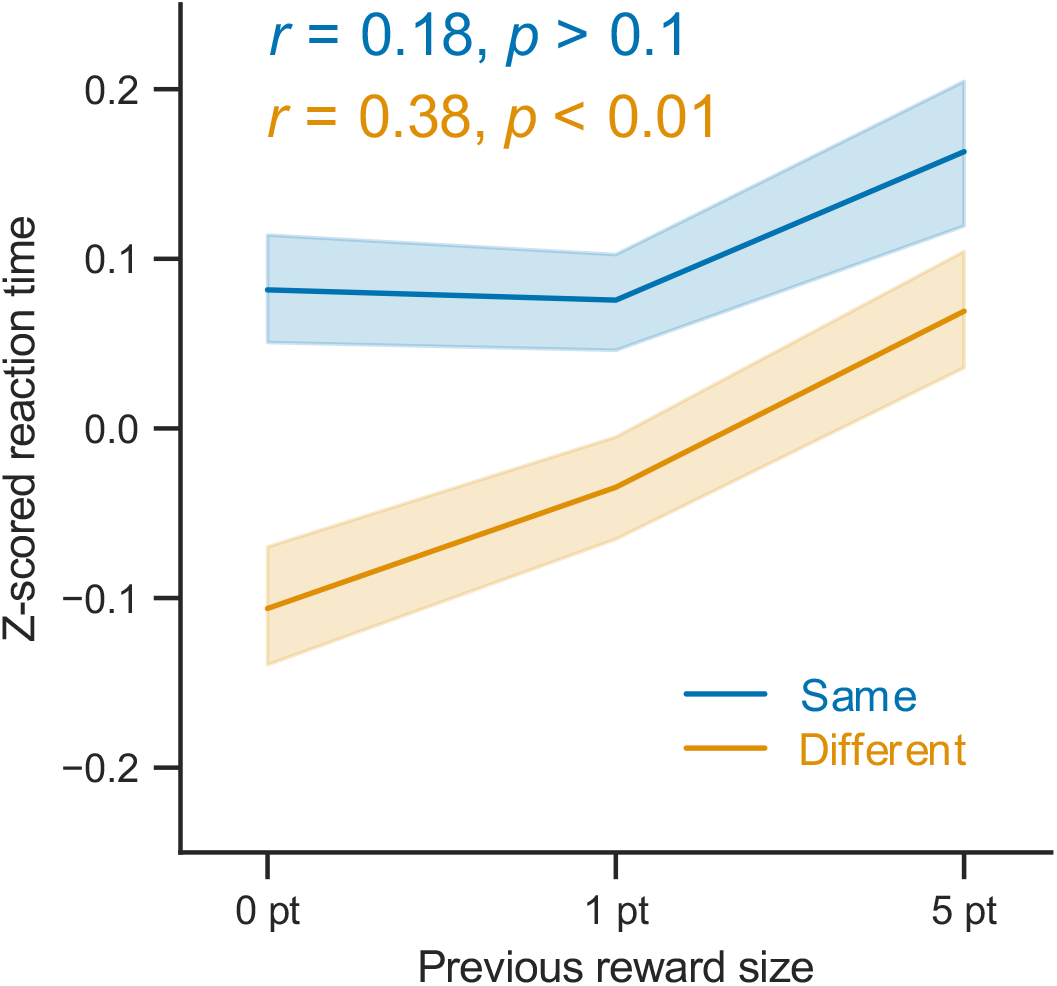
Mean z-scored RT in previous reward size. Each color corresponds to previous-current stimulus condition.

## 4 DISCUSSION

This study demonstrated that reward priming distorted the decision process even when participants were informed explicitly that rewards were irrelevant to the task. In particular, reward priming slowed their responses when the stimulus was the same as in the one in the previous trial but did not accelerate such when the stimulus was different from that in the previous trial (Fig. 7). This evidence is partially consistent with extant literature on random rewards [1, 6, 15] and suggests that irrespective of whether participants are informed of reward randomness explicitly or implicitly, reward priming influences behaviors.

Previous studies in visual search task showed that reward priming accelerates response speed with implicit knowledge of rewards [6, 16, 17]. However, our result showed that reward priming did not accelerate response speed when the stimulus was the same as in the one in the previous trial. This discrepancy suggests that reward priming enhancement depends on the knowledge of rewards rather than actual rewards or performance; that is, the facilitation by reward priming is driven by top-down process.

On the contrary, reward priming suppressed the participants’ responses when the stimulus was different from the one in the previous trial. This concurs with extant literature on reward priming [1, 3, 5, 6]. The reward priming inhibition may influence decisions independent of the knowledge of rewards; the inhibition of reward priming is modulated by actual reward size; that is, the suppression by reward priming is driven by bottom-up process. Therefore, there is a possibility that reward priming comprises partially distinct processes. While it is possible that these results are task-dependent, several experimental tasks have shown reward-driven effects [5, 15, 18, 19], suggesting that the effects of reward priming are based on a domain-general mechanism.

## 5 CONCLUSION

This study revealed that the enhancement and inhibition of reward priming could be partially distinct processes owing to the knowledge of rewards. This result suggests the effect of reward priming consists of top-down and bottom-up processes. The findings extend the comprehension of reward priming in relation to explicit knowledge of rewards, auditory domain, and perceptual decisions. It is recommended that future research explore how aspects related to reward priming modulate perceptual decisions.

## ACKNOWLEDGMENT

We would like to express our gratitude toward Yuma Osako for his valuable advice regarding the experimental design.

## REFERENCES

[1] Anderson BA, Laurent PA, Yantis S (2011) Value-driven attentional capture. Proc Natl Acad Sci U S A 108:10367–10371

[2] Anderson BA (2019) Neurobiology of value-driven attention. Curr Opin Psychol 29:27–33

[3] Della Libera C, Chelazzi L (2006) Visual selective attention and the effects of monetary rewards. Psychol Sci 17:222–227

[4] Failing M, Theeuwes J (2018) Selection history: How reward modulates selectivity of visual attention. Psychon Bull Rev 25:514–538

[5] Kim AJ, Lee DS, Anderson BA (2021) Previously reward-associated sounds interfere with goal-directed auditory processing. Q J Exp Psychol 74:1257–1263

[6] Hickey C, Chelazzi L, Theeuwes J (2010) Reward changes salience in human vision via the anterior cingulate. J Neurosci 30:11096–11103

[7] Sali AW, Anderson BA, Yantis S (2014) The role of reward prediction in the control of attention. J Exp Psychol Hum Percept Perform 40:1654–1664

[8] Deci EL (1972) The effects of contingent and noncontingent rewards and controls on intrinsic motivation. Organ Behav Hum Perform 8:217–229

[9] Della Libera C, Perlato A, Chelazzi L (2011) Dissociable effects of reward on attentional learning: from passive associations to active monitoring. PLoS One 6:e19460

[10] Fröber K, Dreisbach G (2014) The differential influences of positive affect, random reward, and performance-contingent reward on cognitive control. Cogn Affect Behav Neurosci 14:530–547

[11] Stürmer B, Nigbur R, Schacht A, Sommer W (2011) Reward and punishment effects on error processing and conflict control. Front Psychol 2:335

[12] Sohoglu E, Chait M (2016) Neural dynamics of change detection in crowded acoustic scenes. Neuroimage 126:164–172

[13] Vallat R (2018) Pingouin: statistics in Python. J Open Source Softw 3:1026

[14] Holland MK, Lockhead GR (1968) Sequential effects in absolute judgments of loudness. Percept Psychophys 3:409–414

[15] Infanti E, Hickey C, Menghi N, Turatto M (2017) Reward-priming impacts visual working memory maintenance: Evidence from human electrophysiology. Vis Cogn 25:956–971

[16] Hickey C, van Zoest W (2012) Reward creates oculomotor salience. Curr Biol 22:R219–20

[17] Hickey C, Los SA (2015) Reward priming of temporal preparation. Vis Cogn 23:25–40

[18] Anderson BA (2016) Value-driven attentional capture in the auditory domain. Atten Percept Psychophys 78:242–250

[19] Hickey C, Peelen MV (2017) Reward Selectively Modulates the Lingering Neural Representation of Recently Attended Objects in Natural Scenes. J Neurosci 37:7297–7304

